# GeneCompass: Deciphering Universal Gene Regulatory Mechanisms with Knowledge-Informed Cross-Species Foundation Model

**DOI:** 10.1101/2023.09.26.559542

**Authors:** Xiaodong Yang, Guole Liu, Guihai Feng, Dechao Bu, Pengfei Wang, Jie Jiang, Shubai Chen, Qinmeng Yang, Yiyang Zhang, Zhenpeng Man, Zhongming Liang, Zichen Wang, Yaning Li, Zheng Li, Yana Liu, Yao Tian, Ao Li, Jingxi Dong, Zhilong Hu, Chen Fang, Hefan Miao, Lina Cui, Zixu Deng, Haiping Jiang, Wentao Cui, Jiahao Zhang, Zhaohui Yang, Handong Li, Xingjian He, Liqun Zhong, Jiaheng Zhou, Zijian Wang, Qingqing Long, Ping Xu, The X-Compass Consortium, Hongmei Wang, Zhen Meng, Xuezhi Wang, Yangang Wang, Yong Wang, Shihua Zhang, Jingtao Guo, Yi Zhao, Yuanchun Zhou, Fei Li, Jing Liu, Yiqiang Chen, Ge Yang, Xin Li

**Author notes:** These authors contributed equally to this work (Xiaodong Yang, Guole Liu, Guihai Feng, Dechao Bu, Pengfei Wang, Jie Jiang, Shubai Chen, Qinmeng Yang). Correspondence (X. Li), (G. Yang), (Y. Chen), (J. Liu), (F. Li), (Y. Zhou), (Y. Zhao), (J. Guo), (S. Zhang), (Y. Wang), (G. Feng). A list of affiliations appears at the end of the paper (The X-Compass Consortium).

## Abstract

Deciphering the universal gene regulatory mechanisms in diverse organisms holds great potential to advance our knowledge of fundamental life process and facilitate research on clinical applications. However, the traditional research paradigm primarily focuses on individual model organisms, resulting in limited collection and integration of complex features on various cell types across species. Recent breakthroughs in single-cell sequencing and advancements in deep learning techniques present an unprecedented opportunity to tackle this challenge. In this study, we developed GeneCompass, the first knowledge-informed, cross-species foundation model pre-trained on an extensive dataset of over 120 million single-cell transcriptomes from human and mouse. During pre-training, GeneCompass effectively integrates four types of biological prior knowledge to enhance the understanding of gene regulatory mechanisms in a self-supervised manner. Fine-tuning towards multiple downstream tasks, GeneCompass outperforms competing state-of-the-art models in multiple tasks on single species and unlocks new realms of cross-species biological investigation. Overall, GeneCompass marks a milestone in advancing knowledge of universal gene regulatory mechanisms and accelerating the discovery of key cell fate regulators and candidate targets for drug development.

## Introduction

Vertebrate organisms are intricate systems composed of up to trillions of cells that classified into hundreds of different types. These cells collaborate to form different tissues and organs, each serving a unique set of physiological functions^1,2^. Elucidating the gene regulatory mechanisms underlying these tissues and organs is crucial for deciphering the patterns of individual development and promoting clinical therapies. With the rapid advances in omics sequencing technologies, we have now started to dissect how cells in various organs exert their specific functions in single cell resolution^3^ and accumulated large amount of single-cell data. However, gene expression is regulated at multiple levels, ranging from chromatin accessibility to post-transcriptional modification^4,5^. This implies that comprehensively deciphering the gene regulatory mechanisms solely through wet biological experiments is labor-intensive and time-consuming. The emergence of deep learning models, with their ability to capture and represent complex patterns in large datasets, offer an opportunity to dissect the multi-level and cross-species regulatory mechanisms^6,7^.

In recent years, foundation models such as BERT^8^, GPT^9,10^, PaLM^11,12^, and LLaMA^13^ in natural language domains, along with DALL-E^14^ in visual domains have achieved remarkable performance across a wide variety of downstream tasks. They typically adopt a paradigm of initial pre-training on extensive data through self-supervised learning, followed by an adaptation step for specific downstream tasks through fine-tuning. Similar to natural language serving as an abstract layer for understanding human activities, the transcriptome serves as a representative layer for understanding gene regulatory activities within biological systems. Several studies have utilized single-cell transcriptomic data to construct pre-trained foundation models, such as scGPT^15^, Geneformer^16^, and scFoundation^17^. These works share the commonality of leveraging tens of millions of human single-cell transcriptomic profiles to pre-train foundation models and have demonstrated remarkable performance across a broad range of downstream tasks, such as cell clustering, cell type annotation, gene perturbation simulation, and drug target prediction. However, current models have their limitations. Given that vertebrate organisms exhibit vast diversity across species, and the evolutionary conservation of gene regulatory mechanisms, the integration of datasets from different species represents a remarkable opportunity to unravel the intricate complexities of gene regulation^2,18–20^, while current models solely rely on data from a single species. Additionally, the wealth of biological knowledge accumulated over previous decades, encompassing core regulatory region data, experimentally confirmed gene interactions, and gene family annotations, represents our comprehensive understanding of biological processes to date. Infusing these knowledges into the pre-training process can substantially guide the model learn the universal gene regulatory mechanism in a self-supervised manner.

In this study, we proposed GeneCompass, a knowledge-informed cross-species foundation model pre-trained on scCompass-126M, a currently largest corpus encompassing over 120 million single-cell transcriptomes from both humans and mice to our knowledge. Importantly, the model incorporates prior biological knowledge including promoter sequences, gene co-expression networks, gene family information, and transcription factor-target gene regulatory relationships. Through fine-tuning our pre-trained model on various downstream tasks, we have achieved superior or comparable performance to the state-of-the-art (SOTA) models across diverse biological context. Overall, our model represents a significant breakthrough in the development of foundation models for dissecting universal gene regulatory mechanisms across species and expediting the identification of crucial regulators of cell fate and potential targets for drug development.

## Results

### GeneCompass architecture and pre-training

GeneCompass is a knowledge-informed cross-species foundation model pre-trained on transcriptomic corpus of over 120 million cells from human and mouse. GeneCompass could be efficiently applied in various downstream biological-related tasks by finetuning with limited data (Fig. 1a). Utilizing the self-attention mechanism for explicit context encoding^21^, GeneCompass could understand the essence of cells and the intricate relationships among genes based on the input transcriptome.

**Fig. 1.**
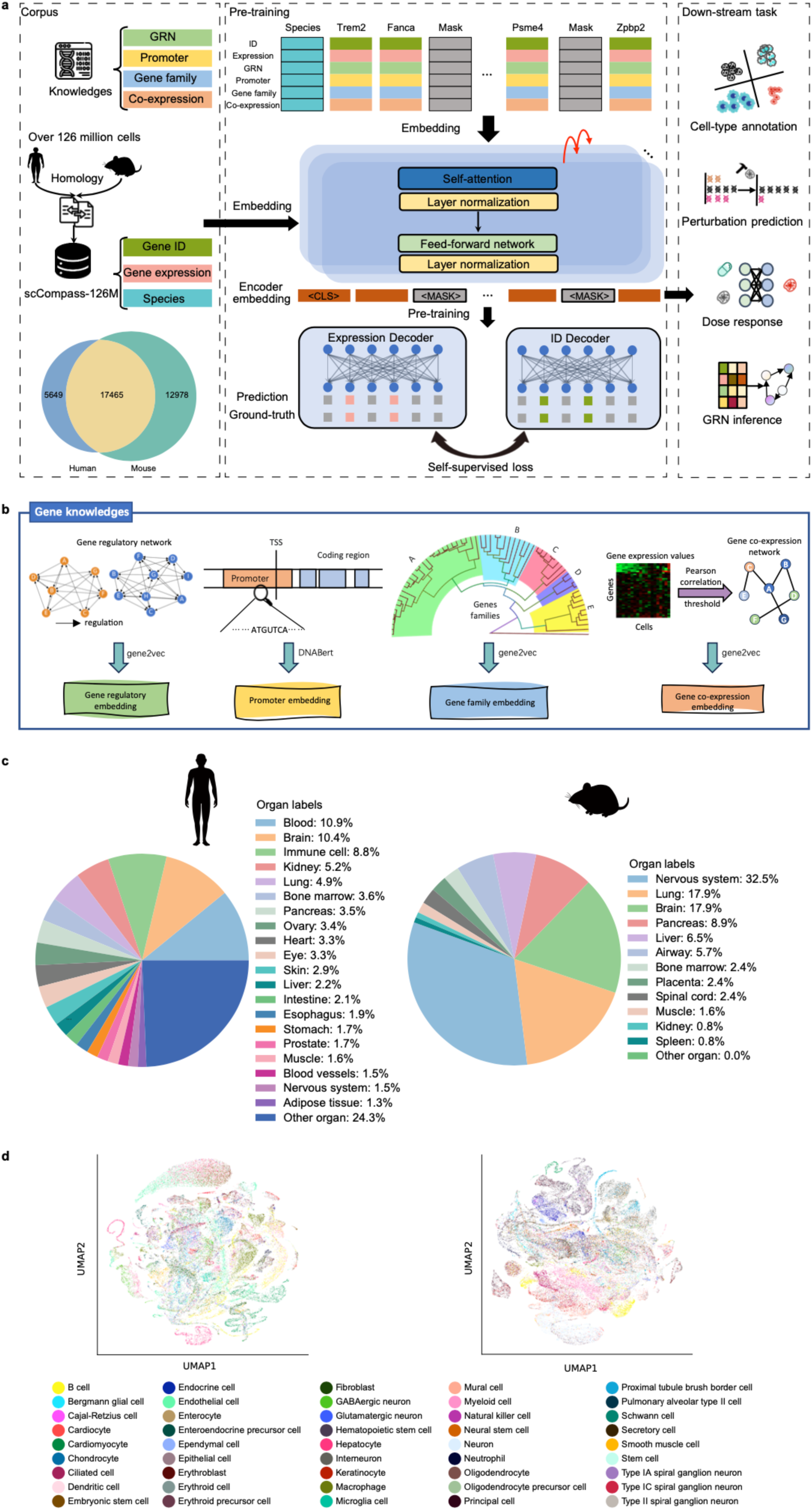
GeneCompass architecture and pre-training corpus. **a,** The framework of GeneCompass. The model is pre-trained on large-scale single-cell transcriptomes from human and mouse. Rather than training from scratch, the pre-trained GeneCompass is used on multiple downstream tasks including cell-type annotation, gene regulatory network prediction, dosage response prediction, etc. **b,** Four prior knowledge embeddings, including gene regulatory network, promoter sequence, gene family and co-expression. **c,** Organ types of human and mouse in scCompass-126M. **d,** UMAP of different cell types of a sampled subset from scCompass-126M.

We initiated the development of GeneCompass by constructing a large-scale pre-training corpus, scCompass-126M. This corpus consists of 126 million single-cell transcriptomes from human and mouse, which is collected from publicly available datasets including an extensive range of organs and cell types (Fig. 1c, d). To ensure data quality, we filtered out cells with outlier gene expression. Additionally, we retained the informative genes with sufficient variability or expression levels across the dataset to capture biological heterogeneity and cell-type-specific signatures. Since there exists a different gene list between human and mouse, we aligned their homologous genes by sharing the same gene IDs. In this study, the token dictionary contains 17,465 homologous genes out of total 36,092 genes (Fig. 1a).

The current landscape of large-scale transcriptomic pre-training models primarily utilize relative gene rankings^16^ or binned gene expression values^15^ as input, leading to inadequate representations of transcriptome. To overcome this limitation, absolute gene expression values and corresponding gene IDs are concatenated and then ranked within the cell based on the normalized expression values across the entire scCompass-126M. (Fig. 1a). The introduction of absolute expression values contributes more fine-grained information and leading to stronger supervision constraints in self-supervised learning to GeneCompass. To further enhance the capability of our pre-trained model, four different types of biological prior knowledge, including promoter sequences, gene co-expression networks, gene family, and transcription factor-target regulation networks, are encoded into a unified embedding space by different methods^22,23^ (Fig. 1b). To denote the species information, a species token embedding is prepended to each cell (Fig. 1a). Overall, GeneCompass integrates gene IDs, expression values and prior knowledge together as gene tokens, and utilizes a 12-layer transformer framework^8,21^ for cell encoding.

Inspired by self-supervised learning in natural language processing (NLP) domain, the masked language modeling (MLM) strategy^8^ is employed to randomly mask gene tokens including gene IDs, expression values and prior knowledge during the pre-training. To be detailed, 15% genes are masked randomly for each cell, and then GeneCompass recovers gene IDs and expression values of those masked gene simultaneously through two separate decoding heads. This multi-task self-supervised learning paradigm, which combines the recovery of relative ranks and absolute values, enhances the ability to capture the intricate and nuanced complexities of gene expression and cell states.

### GeneCompass is capable of species integration and in silico gene network analysis

Homologous genes often retain similar expression patterns and functional roles. This makes the utilization of homology information an effective approach for integrating corpus across species^24^. We hypothesize that homologous genes tend to behave similarly across species, while each different genes behaves variously, reflecting the conservation and diversity of genes. To validate whether the gene embeddings encoded by GeneCompass retain the homology information, we compared the cosine similarity of gene embeddings between different genes within the same species, homologous and non-homologous genes across different species. 2,000 B cells are randomly selected from human and mouse corpus, respectively. In consistent with our hypothesis, homologous genes in GeneCompass are more similar than non-homologous ones between human and mouse in terms of the statistical distribution of the gene embedding similarity. While distinct genes shows distinguishable embedding similarity as expected regardless of whether the cell is human or mouse origin (Fig. 2a). These results indicates GeneCompass successfully captures the conserved patterns and diversity of genes.

**Fig. 2.**
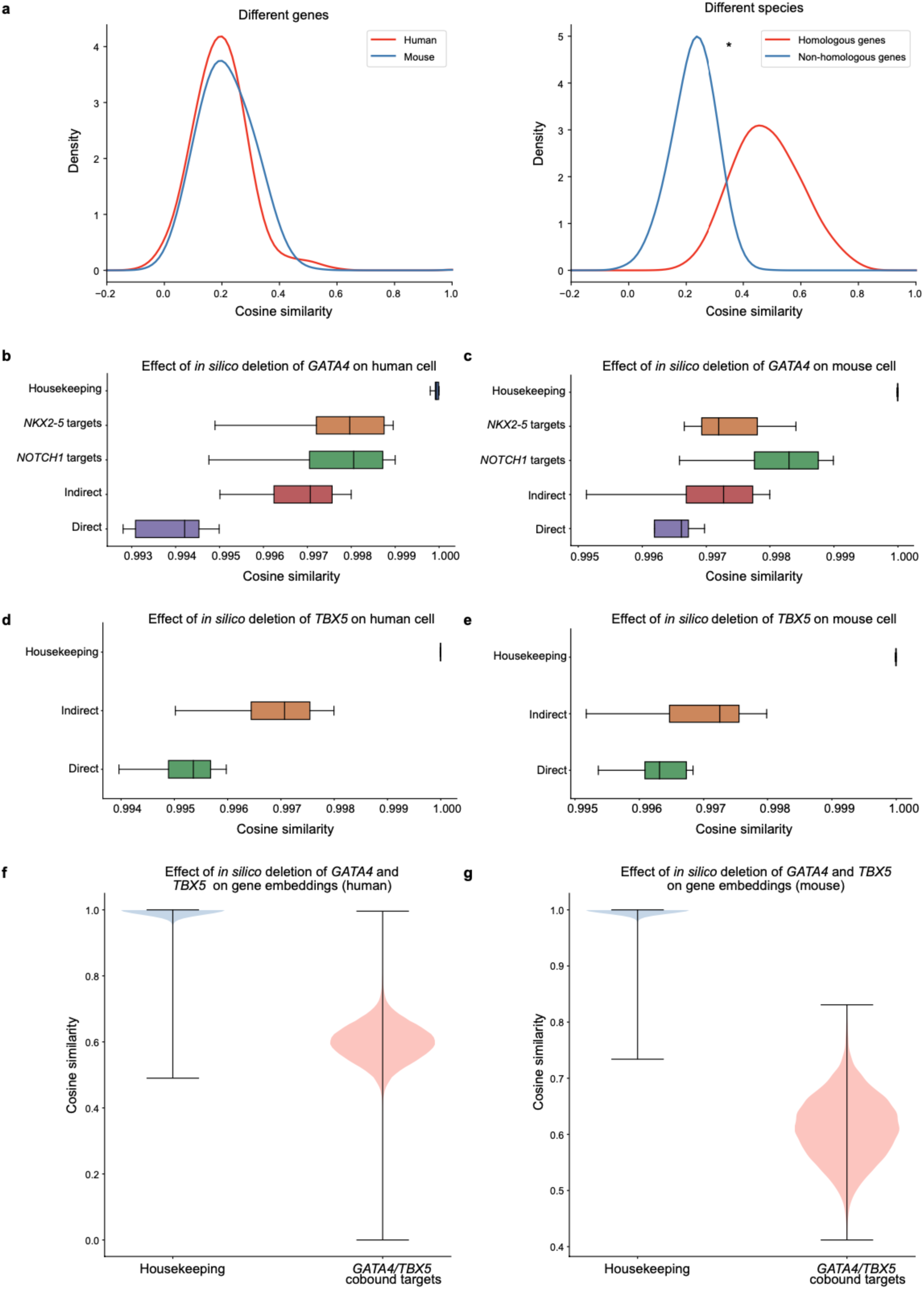
Analysis of gene embedding. **a,** Cosine similarity between different genes of the same species and non-homologous genes of different species (*P < 0.05 Wilcoxon, FDR-corrected). **b-c,** Effects of i*n silico* deletion of *GATA4* on different gene types, including *GATA4* direct targets, housekeeping genes, *NOTCH1* targets, *NKX2-5* targets and *GATA4* indirect targets in human and mouse, respectively. **d-e,** Effects of i*n silico* deletion of *TBX5* on different gene types, including *TBX5* direct targets, *TBX5* indirect targets, and housekeeping genes in human and mouse, respectively. **f-g,** Effects of the combined deletion of *GATA4* and *TBX5* on cobound target genes and housekeeping genes in human and mouse, respectively.

In consideration that GeneCompass have captured the gene interaction mechanism via the pre-training process, we assume *in silico* gene perturbation assay could simulate the biological process at gene level. In previously reports, the deletion of congenital heart disease related genes, *GATA4* and its binding partner *TBX5*, had a significantly greater effect on their direct target genes compared with non-directed target genes, which was observed in fetal cardiomyocytes^25,26^. We compare the cosine similarity difference of a given gene list when *in silico* deleting *GATA4* and *TBX5* individually or simultaneously in human cells. Results show that deleting *GATA4* or *TBX5* individually has a significantly more deleterious impact on their direct targets than on indirect targets and housekeeping genes (Fig. 2b-e). The result also suggests that the computational approach recognizes the cooperative action of *GATA4* and *TBX5* at these cobound targets (Fig. 2f-g). These findings are consistent with those in Geneformer. Additionally, as GeneCompass integrates the homologous information across species, we conduct the *in silico* perturbation of GATA4 and TBX5 in mouse cells, yielding similar results with human (Fig. 2b-e). Therefore, the extensive experiments validate the homology and the gene interaction information are recognized in the gene embeddings encoded by GeneCompass, which would promote both zero-shot and fine-tuning downstream tasks.

### GeneCompass boosts cell-type annotation from single-species to cross-species

While existing methods have shown decent performance on cell type annotation, they only focus on single-species tasks. GeneCompass is pre-trained by the cross-species corpus, which significantly boosts cell-type annotation from single-species to cross-species. For single-species cell type annotation, GeneCompass outperforms alternative methods when using the same amount of task-specific data for fine-tuning (Fig. 3a). With the same size of corpus, GeneCompass is consistently better than other benchmarked methods such as Geneformer and scGPT in terms of accuracy and macro-f1. Additionally, on the task of human-specific cell type annotation, GeneCompass pre-trained with a combined human and mouse corpus demonstrates superior performance compared to models trained solely on the human data, or models trained on equivalent amounts of mouse data. This indicates that adding the corpus of different species from the downstream task to pre-training could further improve the fine-tuned performance.

**Fig. 3.**
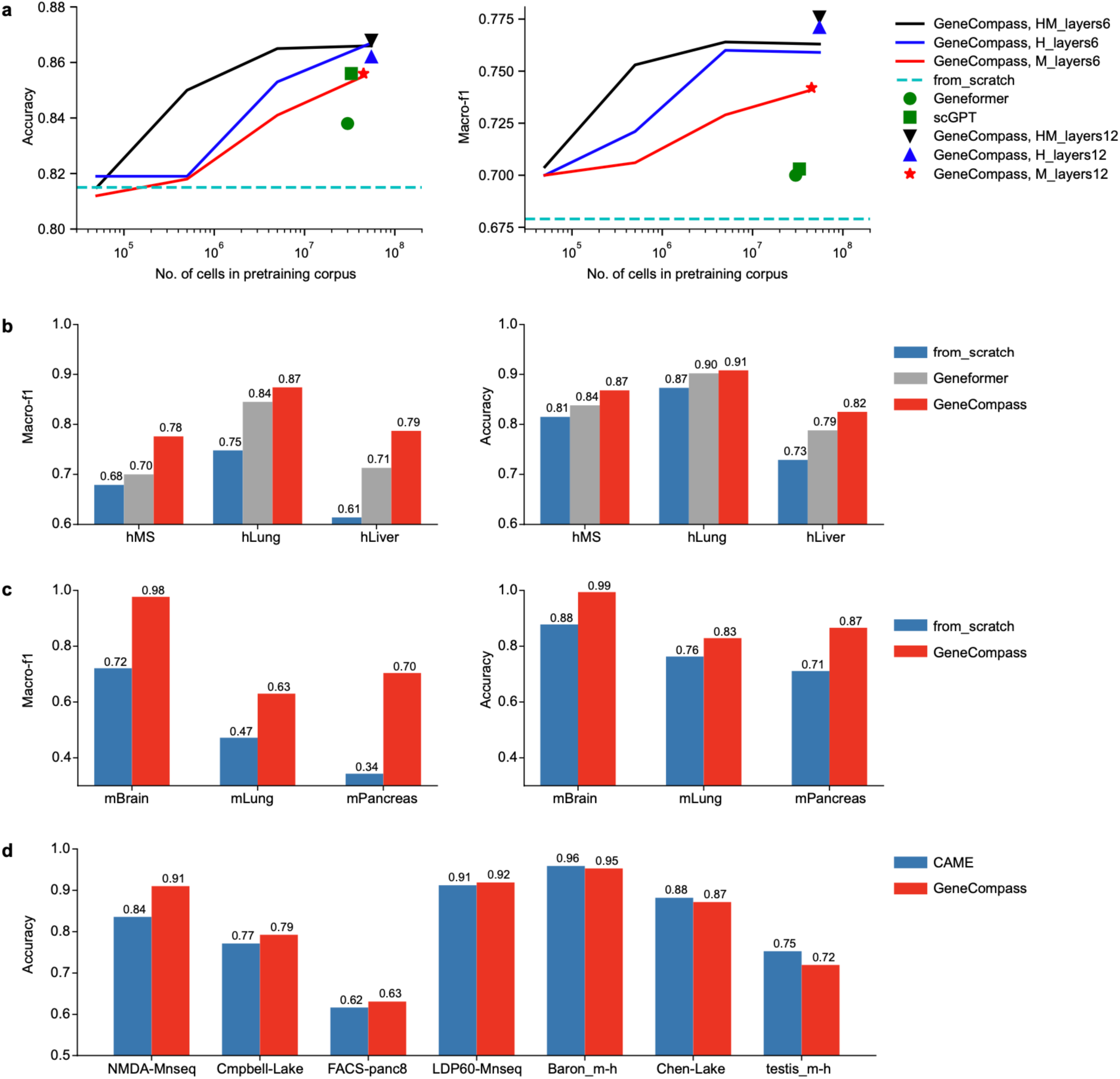
GeneCompass boosts the performance of cell type annotations from single-species to cross-species. **a,** Performance comparison of GeneCompass and other benchmarked methods on the downstream task of cell type annotation on human multiple sclerosis (hMS) dataset. GeneCompass is pre-trained by the human-mouse (HM, black line), human (H, blue line), and mouse (M, red line) single-cell transcriptomes corpus with different cell numbers, respectively. “layers6” and “layers12” denote self-attention transformer with 6 layers and 12 layers, respectively. The cyan dashed line represents the result of randomly initialized GeneCompass. The green circle point and green square point represent the results of Geneformer and scGPT, respectively. **b,** Performance of GeneCompass on hMS, hLung, and hLiver datasets. **c,** Performance of GeneCompass on mBrain, mLung, and mPancreas datasets. **d,** Cross-species cell type annotation from mouse to human of GeneCompass. The datasets in **b-d** derived from human and mouse marked as "h" and "m" respectively and the detailed information of datasets can be found in Supplementary Materials.

To validate the effectiveness of biological prior knowledge integration, an ablation study is performed on the task of human cell annotation (Extended Fig. 1a). The results show that the infusion of gene regulatory network (GRN), co-expression (co-exp), or gene-family embedding individually leads to a significant improvement in accuracy. While the infusion of promoter embedding solely does not enhance cell annotation performance, we anticipate potential improvements in other tasks. Notably, when all four knowledge embeddings are combined, there is also an enhancement in cell annotation performance.

In order to evaluate the performance of GeneCompass in a rigorous way, we perform single-species cell-type annotation analysis on datasets of diverse organs from human and mouse. GeneCompass always achieves the best performance on all datasets. Compared with training from scratch, GeneCompass improves the macro-f1 by 10%, 12% and 18% on different human datasets (i.e., hMS, hLung and hLiver) and by 26%, 16% and 36% on different mouse datasets (i.e., mBrain, mLung and mPancreas), respectively. Furthermore, we compare our model with the SOTA methods. Compared with Geneformer, GeneCompass improves the macro-f1 by 3∼8% on human datasets. Compared with TOSICA^27^, a recent transformer-based cell type annotation method, GeneCompass achieves higher recall on 16 cell types of all 18 types on the mPancreas dataset (Extended Fig.1b-d). These results indicate the superiority of GeneCompass on multiple-species cell type annotation tasks by using the large-scale cross-species corpus and prior knowledge infusion.

To explore the capability of GeneCompass on cross-species downstream tasks, we integrated GeneCompass with the SOTA method CAME^28^ for cross-species cell type annotation. Gene embeddings generated by GeneCompass are utilized as initial gene-nodes features within CAME. We embarked on a comprehensive evaluation on seven paired datasets sourced from four distinct organs (i.e., brain, pancreas, retina, and testis). By leveraging the cell type annotations of the mouse, we ventured into the realm of forecasting human cell types. Each dataset pair underwent twenty distinct experiments, and their averages served as the yardstick for comparison. Following the integration of GeneCompass, a subtle yet noteworthy improvement is shown in the prediction accuracy in comparison to CAME (Fig. 3d). Particularly, GeneCompass shows a pronounced improvement in specific context such as retina, with a surge of 7.5% in accuracy. This demonstrates that GeneCompass, through pre-training on cross-species data, has generated gene embeddings with the capability to empower cross-species predictions.

### Pre-trained gene embedding elevates GRN inference, drug response and gene dosage sensitivity prediction tasks

To further validate the capability of gene embeddings encoded in GeneCompass, we investigate several downstream tasks including GRN inference, drug response and gene dosage sensitivity. The GRN provides information on gene regulation and signal transduction, offering insights into gene expression patterns and key regulatory genes in diseases. To evaluate the benefit of pre-trained models on GRN inference, we apply gene embeddings of each model on DeepSEM^29^, an advanced GRN inference tool. The AUPRC (area under the precision-recall curve) of the GeneCompass model increases consistently with increasing data volume. Compared with other competing methods, GeneCompass trained by 55million human data has the best performance in GRN inference (Fig. 4a).

**Fig. 4.**
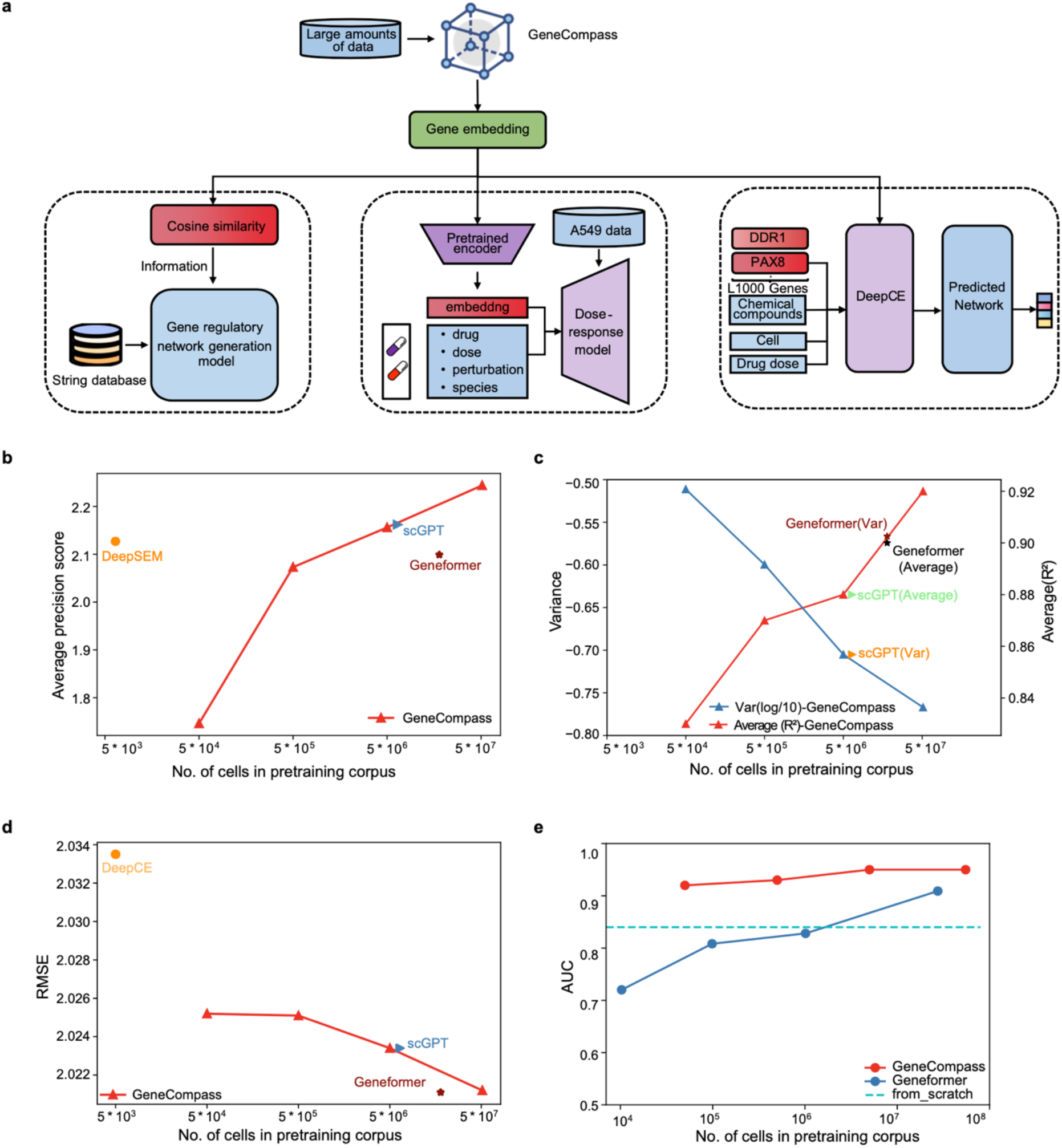
GeneCompass elevates GRN inference, drug response, gene expression profile, and gene dosage sensitivity prediction tasks. **a,** The framework of three downstream tasks based on gene embeddings of GeneCompass. **b,** Performance comparison on the GRN inference task. **c**, Performance comparison on the dose response task. **d,** Performance comparison on the gene expression profile prediction task. **e,** Performance comparison on the dosage sensitivity transcription factors prediction task.

Predicting cellular responses to different types of drugs and their dosages is key to promote drug development. We next test GeneCompass on drug dose response prediction. Compositional Perturbation Autoencoder (CPA) model is applied on this task^30^. Similar with GRN, gene embeddings produced from each pre-trained model are added into CPA. We calculate the average R-Squared score and variance of results from each model. Results show that with the increase of data volume, GeneCompass exhibits a constantly climbing trend in performance. Notably, GeneCompass pre-trained by 55 million of human data achieves the highest score (Fig. 4b). Moreover, GeneCompass shows lower variance for different drug conditions compared with Geneformer (Extended Fig.2d).

Gene expression profiling helps describe phenotypic features and advances drug discovery. DeepCE is a widely used model in the field of predicting drug-induced gene expression perturbations^31^. We are attempting to incorporate gene embedding information generated by GeneCompass into DeepCE to assess its impact on the model’s performance. We calculate root mean squared error (RMSE) of results from each model. As the data volume increases, GeneCompass continues to exhibit a consistently improving trend in performance (Fig. 4d).

In the field of genetic diagnosis, determining which genes are dosage-sensitive poses a significant challenge when interpreting copy number variations (CNVs). To address this, we utilize pre-defined dosage-sensitive and non-sensitive gene sets as training data and employ GeneCompass’s gene embedding to construct a fine-tuned model for dosage sensitivity gene identification. We observe that as the number of cells used for pre-training increases, the predictive performance of GeneCompass in the gene dosage sensitivity prediction task consistently improves (Fig. 4e). Specifically, the area under the receiver operating characteristic curve (AUC) reaches up to 0.95. It is worth mentioning that the gene embedding from GeneCompass outperforms Geneformer across all training data sizes in this task.

### Pre-trained gene embedding elevates gene perturbation prediction

Compared with the impact of drug treatment or changes in gene copy numbers on gene expression dosage, functional mutations in genes play a prominent role in causing diseases in clinical practice. GeneCompass learns complex interactions between genes and incorporates this knowledge into their gene embeddings. We attempt to leverage the gene embedding provided by GeneCompass to predict cellular changes resulting from perturbations caused by functional gene mutations. We integrate the gene embedding of GeneCompass into the advanced perturbation prediction tool GEARS, which learns a gene embedding from a gene co-expression knowledge graph and a perturbation embedding from a Gene Ontology (GO)-derived knowledge graph^32^. We replaced the original gene embeddings with those from GeneCompass (Fig. 5a). This leads to a 15.4% reduction in Mean Squared Error (MSE) for the top 20 differentially expressed (DE) genes, indicating a decrease in prediction bias for these critical genes (Fig. 5b).

**Fig. 5.**
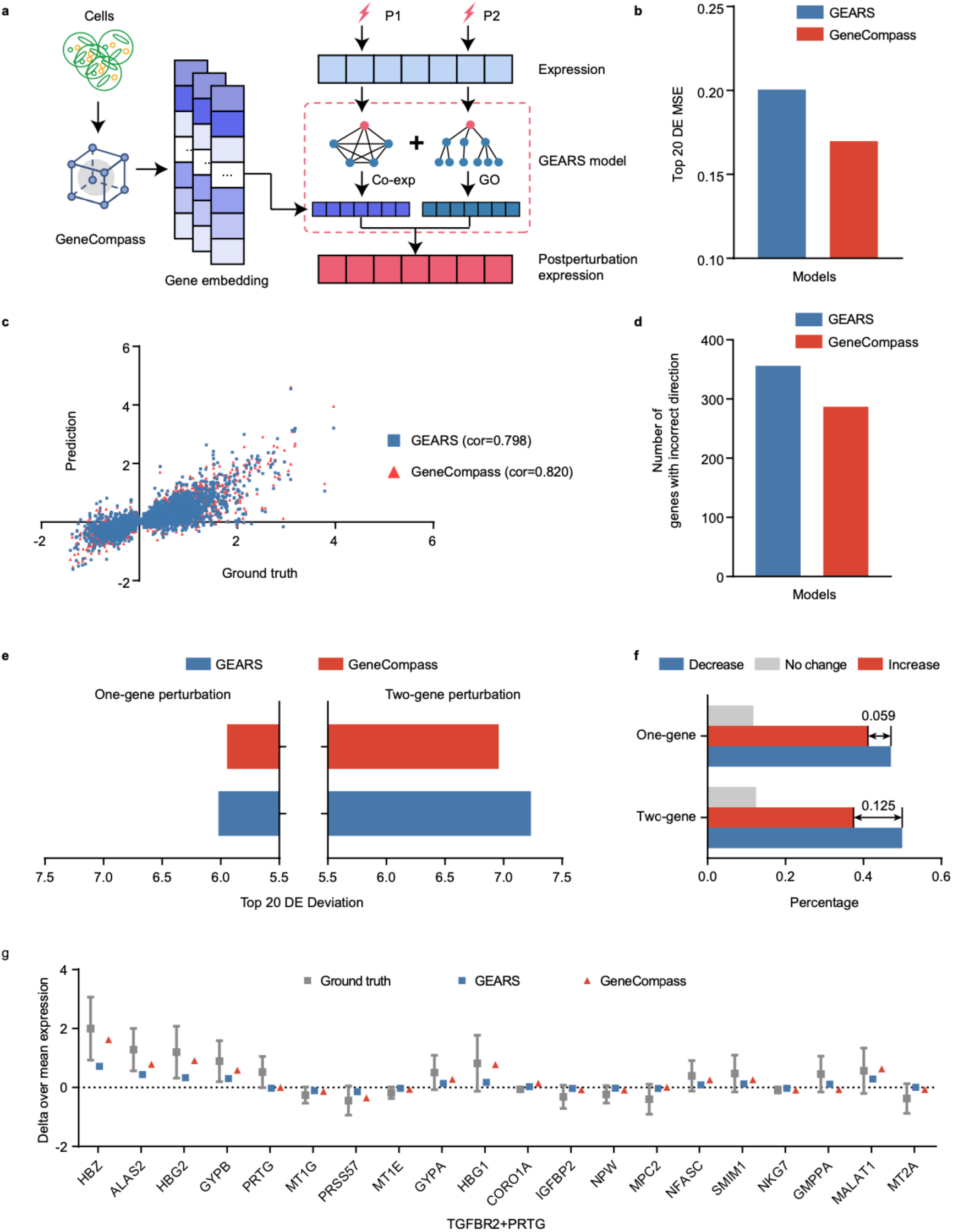
GeneCompass elevates gene perturbation prediction task. **a,** The workflow of GeneCompass for perturbation prediction task. We replaced the GEARS’ gene embedding learned from gene co-expression knowledge graph with GeneCompass’ gene embedding. **b,** Mean Square Error (MSE) in predicting top 20 differentially expressed (DE) genes’ expression change by GeneCompass and GEARS. MSE was measured on 20 most DE genes between true postperturbation expression and prediction results. **c,** Scatter plot of the predicted postperturbation gene expression change and true postperturbation expression change. Each dot represents a specific gene, and the Pearson correlation coefficient is marked as "cor". **d,** Total number of the top 20 DE genes where the predicted postperturbation differential expression in the incorrect direction of the ground truth. **e,** Expression deviation between the predicted postperturbation gene expression and true postperturbation expression for top 20 DE genes by GeneCompass and GEARS. **f,** Percentage of prediction results by GeneCompass showing a decrease, no change, or increase in the top 20 differential expression (DE) deviation for one-gene and two-gene perturbations compared to GEARS. **g,** Expression change for the combined *TGFBR2* and *PRTG* perturbation in true experiment postperturbation, predicted by GeneCompass and GEARS. Box-whiskers indicate mean gene expression with standard deviation (SD) after perturbing the gene combination *TGFBR2* and *PRTG* (n=205). The red triangle symbol shows the gene expression change predicted by GeneCompass with *TGFBR2* and *PRTG* perturbation excluded during training. The blue square symbol shows the gene expression change predicted by GEARS.

Pearson correlation coefficient (cor) between the predicted post-perturbation gene expression and true post-perturbation expression shows that GeneCompass achieves an improvement of 2.2% compared to GEARS, increasing from 79.8% to 82.0% (Fig. 5c). Next, we investigate whether GeneCompass could more accurately capture the right direction of changes in gene expression following perturbations and summarize the number of top 20 DE genes with incorrect direction for each perturbation prediction. This number decreases by 19.4%, from 356 to 287, compared with results predicted by GEARS (Fig. 5d).The analysis of deviation for each perturbation prediction in the held-out testing reveals that both one-gene and two-gene perturbations exhibit lower summarized top 20 DE deviation in GeneCompass than in GEARS (Fig. 5e). To be specific, GeneCompass provides an extra enhancement of 5.9% for one-gene perturbations and of 12.5% for two-gene perturbations than GEARS (Fig 5f).

Furthermore, we present an example of perturbing the two-gene combination of TGFBR2 and PTRG. Among the top 20 DE genes, the prediction results of 17 genes from GeneCompass are closer to the ground truth compared with results from GEARS (Fig. 5g). In summary, the gene embedding of GeneCompass provides a more effective representation of the relationship between genes, enhancing the prediction of gene perturbations.

### GeneCompass is capable of *in silico* quantitative perturbation for cell reprogramming and differentiation

Owing to pre-training with both ranked gene IDs and expression value, GeneCompass learns the gene-gene regulatory mechanism in an exquisitely fine resolution, which enables quantitative deciphering gene functions. This implies great potential in prediction cell fate transition in different biological process (Fig. 6a). To determine the capacity of *in silico* quantitative perturbation of GeneCompass, we aim to reproduce iPSCs induction by OSKM genes (*Oct4*, *Sox2*, *Klf4* and *c-Myc*^33^), a well-characterized reprogramming paradigm. Two levels of overexpression on OSKM genes are set by the median gene values (low-level overexpress) and the maximum value (high-level overexpress). We also randomly overexpress another four genes as control. In contrast to the control group, the cell state shifts to iPSCs state after *in silico* overexpressing OSKM genes from the initial human fibroblast state in both conditions. A consistent result can be obtained from mouse cells (Fig. 6 b). Besides, the high-level overexpression obtains a higher similarity with iPSCs compared with low-level overexpression. Furthermore, we test GeneCompass on *in silico* quantitative knock-out assay. *Zbtb11* and *Zfp131* are transcription factors (TFs) essential for the pluripotency of Embryonic stem cells (ESCs) for mouse cells^34^. Compared to the original state, *in silico* perturbing ESCs shifts gradually to endoderm status by decreasing their expression of *Zfp131* and *Zbtb11* to the half, the quarter and totally knocked out (Fig. 6c, d). In summary, the results indicate that GeneCompass could simulate the influence of gene expression changes on the cellular fate in a quantitative manner.

**Fig. 6.**
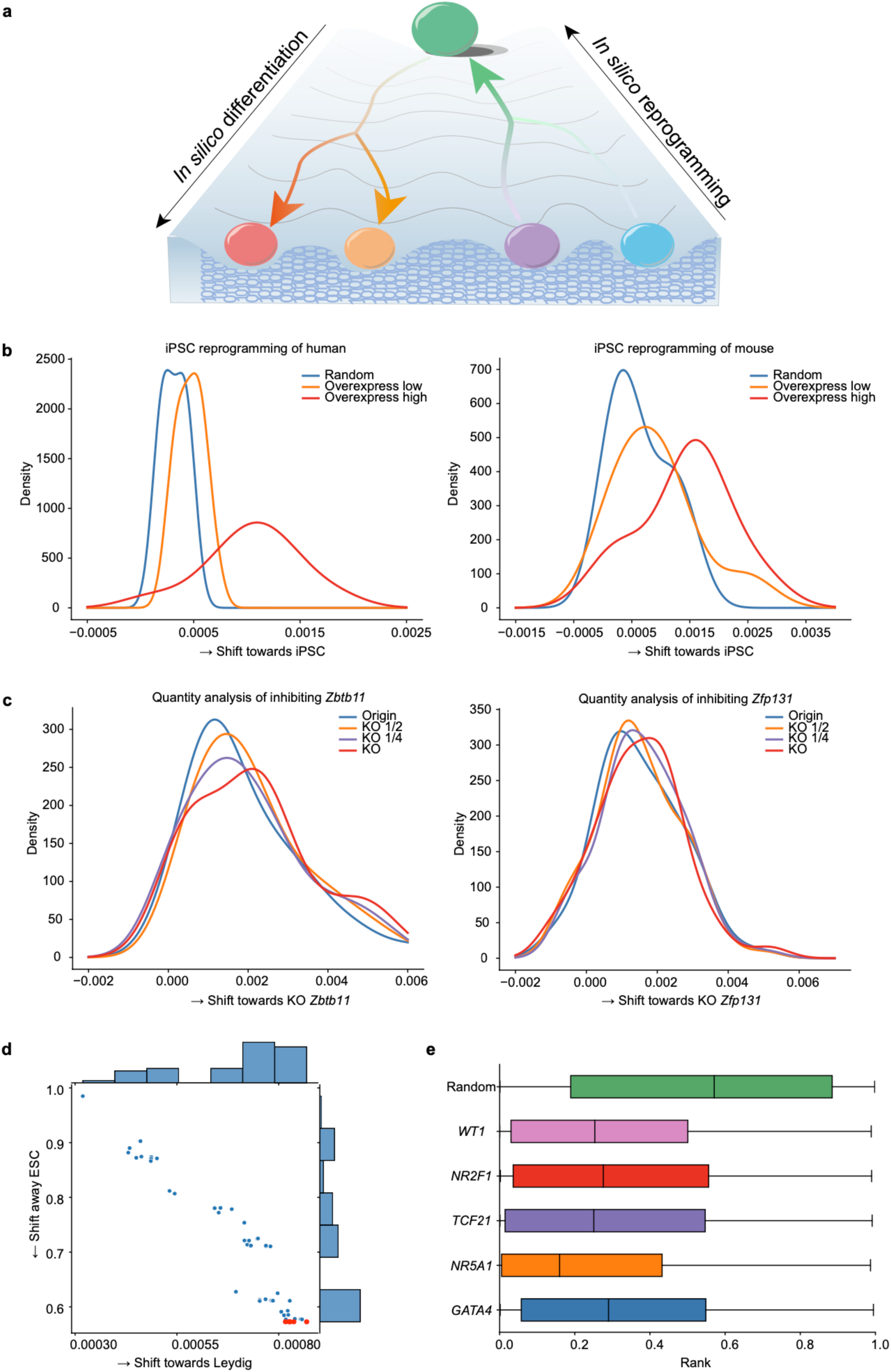
In silico quantitative perturbation for cell reprogramming and differentiation. **a**, Diagram of *in silico* cell fate transition. **b**, *In silico* reprogramming of fibroblasts by artificially overexpressing *OCT4*, *SOX2*, *KLF4*, and *MYC* (OSKM) to the iPSCs state in both human and mouse cells. **c**, *In silico* perturbation of ESCs by artificially decreasing *Zbtb11* or *Zfp131* to the Endoderm state in mouse cells. "KO 1/2" denotes knocking target gene to the half, " KO 1/4" denotes knocking *Zbtb11* or *Zfp131* gene expression to its quarter; and "KO" denotes totally knocking out *Zbtb11* or *Zfp131* gene in the cell. **d**, Distribution of cell embedding shift towards Leydig status in response to *in silico* overexpress of candidate targets in human ESCs cells (top 50 genes towards Leydig status). Selected genes in **e** are marked in red. **e**, By ranking the embedding similarity of Leydig cells and gene-overexpressed ESCs cells, the listed five genes are at the top 30% approximately and set as candidate genes for differentiation.

Considering finding key biological regulators through wet experiments is labor-intensive and time-consuming, it is feasible to apply GeneCompass to facilitate this process. The early gonadal cells are known to be highly plastic and multipotent, which eventually differentiate and give rise to mature and functional somatic cells in adult gonad (testis or ovary)^35^. Systematic single-cell transcriptome analysis of embryonic/fetal gonads in mice and humans has revealed the expression of several key transcription factors that may play critical roles in early gonadal development^36–38^. Here, we perform an *in silico* differentiation analysis with GeneCompass on human ESCs to investigate whether the activation of specific pathways could shift the cell embeddings towards the Leydig state (Fig. 6a). To be specific, each gene in the initial ESCs state is overexpressed to the headmost to get a perturbation state one by one. The perturbation state could reveal the different differentiation directions of ESCs. By comparing the cosine similarities of cell embeddings, we identify five gene regulatory factors including *WT1*, *NR2F1*, *TCF21*, *NR5A1* and *GATA4* exhibiting higher similarity to the target Leydig status while lower similarity to the original ESCs (Fig. 6 d, e). This indicates that these five genes exert significant promotive effects on ESCs differentiation. These findings are also consistent with previously reported studies, demonstrating that *WT1*, *NR5A1* and *GATA4* can facilitate the differentiation of stem cells into Leydig cells.^39–41^ Thus, GeneCompass proves its capability to predict genome-wide screening of genes that promote directed differentiation and is expected to improve the efficiency of wet experiments through the combination of *in silico* (dry lab) and in vitro (wet lab) experiments.

## Discussion

In this study, we introduce GeneCompass, a large-scale pre-trained model that integrates 126 million cross-species single-cell transcriptome and different types of prior biological knowledge. By exploring the embedding information generated by the model for various downstream tasks, GeneCompass shows exceptional performance, surpassing the capabilities of most existing SOTA models. Throughout the model training process, we conduct meticulous ablation experiments to investigate the impact of incorporating various types of prior knowledge on cell type annotation tasks. These experiments prove that prior knowledge has a positive influence on the performance, providing invaluable insights for future developments of foundation models in biology. Furthermore, our study reveals that supplementing the gene expression values alongside the gene ranked information will further expand the model’s capability of quantitative deciphering gene functions in an exquisitely fine resolution.

However, there remains potential for improvement. Currently, our model incorporates information from only two species, i.e., human and mouse. When attempting to include data from other species, we suspect that the introduction of species-specific gene expression patterns may offset the benefits derived from their single-cell data. Nevertheless, cross-species unified gene representation provides valuable insights to establish universal foundation models. Besides, in terms of incorporating prior knowledge, more other essential information such as enhancers and protein sequences has not been explored. Additionally, apart from transcriptional data at the single-cell level, there exists a wealth of epigenomic, proteomic, and metabolomic data that contains rich insights into gene regulation. Investigating effective strategies for integrating this multi-modal information into the model represents a pivotal avenue for future research.

Our current model already demonstrates promising performance in multiple downstream tasks. As it continues to evolve and gain broader adoption, it could offer significant value in optimizing cell fate reprogramming systems, directed organoid cultivation, opening up potential applications in clinical contexts. For instance, it can be applied to discovery of candidate targets of diseases, tumor drug screening and drug toxicity prediction. We anticipate that the ongoing progress and fine-tuning of biological foundation models will bring about significant transformations in both fundamental biological research and clinical applications. In the future, the fusion of dry and wet experiments will create a novel paradigm in life science research, serving as a catalyst for advancements in various areas of the field.

## Methods

### Collecting and preprocessing of multi-species training data

We construct a large-scale pre-training corpus, scCompass-126M. This corpus consists of more than 120 million single-cell transcriptomes consisting of two different species, human and mouse. Multi-species single-cell data provides a rich resource for understanding cellular heterogeneity across diverse organisms. However, collecting and preprocessing such data can be challenging due to differences in biological processes and technical variability between species. Here, we describe the collecting and preprocessing of multi-species single-cell training data from two common model organisms: human and mouse.

Among the species, the cells of human and mouse have the highest ratio, and each of them consists of over 50 million cells. The data used in this study was curated from publicly available datasets collected from various sources, including the National Center for Biotechnology Information (NCBI) Gene Expression Omnibus (GEO), NCBI Sequence Read Archive (SRA), European Molecular Biology Laboratory-European Bioinformatics Institute (EMBL-EBI)-ArrayExpress, and China National Center for Bioinformation (CNCB)-Genome Sequence Archive (GSA). We download FASTQ files from these databases and obtain gene raw counts by running the same pipeline. And other gene raw counts data were downloaded directly from the CELLxGENE, Single Cell Portal, Curated Cancer Cell Atlas (3CA), Cell BLAST, Human Cell Atlas, Temporal Expression during Development Database (TEDD) and some biological studies by other authors^1,18,42,43^. To prepare the multi-species single-cell data for downstream analyses, we perform several preprocessing steps. For quality control, we exclude low-quality and damaged cells, with less than seven genes for proteins or miRNAs. Then we conduct normalization and log1*p* transformation to reduce the skewness.

In short, we collect and preprocess multi-species single-cell training data from human and mouse. The datasets provide a valuable resource for studying cellular heterogeneity across different organisms and for developing machine learning models that can generalize across species. Our approach can be adapted to other multi-species datasets and can facilitate the integration of single-cell data from different organisms for comparative analyses. Single-cell data preprocessing is the systematic procedure of cleaning, standardizing, and refining raw single-cell data into a structured and analyzable format, involving steps such as quality control and gene filtering to remove noise, normalization and transformation to account for technical variations, feature extraction to retain relevant genes.

### GeneCompass architecture and pre-training

#### GeneCompass architecture

GeneCompass employs a 12-layer self-attention transformer^8,21^ to encode input embedding, each composed of a self-attention layer with 12 attention heads, a feed-forward layer and a layer-normalization layer. GeneCompass operates on the sequence of 2048 embedded gene tokens with 768 dimensions, making the number of model parameters reach over 100,000,000. GeneCompass uses bidirectional self-attention based on the number of genes detected in each cell with the pre-trained corpus. During pre-training and finetuning, the Gaussian Error Linear Units (GELUs) are employed as nonlinear activation and the dropout probability for both attention and dense layer is 0.02 (standard deviation of the initializer for weight matrices, 0.02; epsilon for layer normalization layers, 1 × 10^-12^). Code for model configuration, data loading and training is implemented by Pytorch^44^ and Huggingface Transformers library^45^ for model configuration, data loading and training. Furthermore, we extended the library to be capable of inputting scalable external knowledge.

#### GeneCompass pre-training and optimization

GeneCompass integrates the gene ID, expression value and corresponding prior knowledge (promoter, GRN, gene family and co-expression) into together to encode cell transcriptomes. Inspired by self-supervised learning in the Natural Language Processing (NLP) domain, a masked language modeling strategy^8^ is employed to randomly mask genes including their IDs, expressions and prior knowledge during the pre-training. This technique is proven to help large models learn better representations effectively. To be detailed, 15% genes are selected randomly to mask for each cell. Comparing with the existing works, GeneCompass builds a multi-task learning paradigm to predict both the ID and the expression of the masked gene based on the encoded embedding in the meanwhile. We use the Mean Square Error (MSE) objective to optimize the model to predict the gene expression for unknown genes, which is defined as follows:

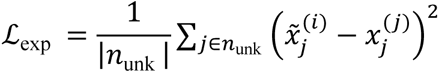

where *n*_unk_ denotes the number of unknown gene expressions and 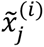 is the predicted gene expression. The ground-truth is denoted as 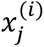.

The cross-entropy loss is employed to optimize the gene ID prediction, which is defined as follows:

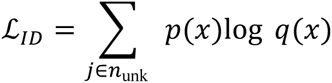

where *p*(*x*) denotes the predictions and *q*(*x*) denotes the real gene token ID.

The pre-training parameters are demonstrated as follows: the learning rate is set to linear decay with 10,000 warm-up steps and the max learning rate is 1e-3 using AdamW optimizer; the batch size is 10 with a gene padding strategy. To make full use of the GPU ability, Deepspeed is applied in the code framework as it provides optimizations that leverage techniques like dynamic batching, mixed precision, model parallelism and memory optimizations to speed up the training of large neural networks while reducing their memory consumption, all within a PyTorch-like API. The pre-training process is accomplished in 9 days using a 4nodes server with eight Nvidia A800 GPUs.

#### Ablation experiment in different data size

An ablation study of GeneCompass using different pre-trained cell numbers is carried out. Specifically, for GeneCompass pre-trained by the human single-cell transcriptomes corpus, different cell numbers (i.e., 5× 10^4^, 5× 10^5^, 5× 10^6^ and 5.5× 10^7^) are used. For GeneCompass pre-trained by the mouse single-cell transcriptomes corpus, different cell numbers (i.e., 5× 10^4^, 5× 10^5^, 5× 10^6^ and 4.5× 10^7^) are used. For GeneCompass pre-trained by the human-mouse single-cell transcriptomes corpus, different cell numbers (i.e., human 5× 10^4^+ mouse 5× 10^4^, human 5× 10^5^+ mouse 5× 10^5^, human 5× 10^6^+ mouse 5× 10^6^, and human 5.5× 10^7^+ mouse 4.5× 10^7^) are used. To make GeneCompass fully converge, larger epochs are used for corpus with smaller cell number (Supplementary Table 2).

#### GeneCompass fine-tuning

The GeneCompass model for finetuning consists of an encoder and a decoder. The encoder has 12 transformer layers which are initialized by the pre-trained weight, and the decoder is the task-specific layer. The finetune model is employed for various downstream tasks such as cell type annotation, dose-response prediction and gene regulation network inference. The fine-tuning hyperparameters for different downstream tasks may be different since the differences in task paradigms. For fair comparison we employ the same hyperparameters (maybe not optimal) in the same task across different datasets, which demonstrates the strong generalization of the GeneCompass.

#### Cell and gene embedding

The 768-dimensions cell embeddings are generated from <CLS> token, which is added to the beginning of single-cell information to represent the characteristics. Besides, GeneCompass could encode each gene to 768 dimensions which contain the context information of a gene in a single cell transcriptome. Note that both cell and gene embedding are obtained from the last layer of transformers.

### Knowledge embedding and incorporation

Four kinds of prior biological knowledge are incorporated into the pre-trained model, including promoter sequences, gene co-expression networks, gene family information, and transcription factor-target gene regulatory relationships.

#### Promoter embeddings

The promoter is the non-coding sequence region of a gene, serving as the activating signal for gene transcription. In our study, the promoter for each gene consists of 2500 bases, including upstream 500 bases before the transcription start site (TSS) and downstream 2000 bases behind TSS. The promoter sequences are finetuned on the pre-trained model DNABert^22^ for 40 epochs to obtain the promoter embeddings with 768 dimensions.

#### Co-expression embeddings

Highly co-expressed gene pairs are genes whose transcriptional expression profiles are highly correlated in a variety of organs or tissues without external interference. Thus, highly co-expressed genes are theoretically related and similar. We hope that after encoding, the distance between co-expressed gene pairs is as close as possible, and the distance between non-co-expressed genes is as far away from each other as possible. This ensures that the similarity between the genes in the high-dimensional space is not affected. Specifically, we calculate the Pearson correlation coefficient (PCC) of each gene pair. Gene pairs with PCC larger than 0.8 are selected for embedding through the gene2vec method^23^.

#### Gene family embeddings

Genes in the same gene family have the same ancestral gene, so the genes are functionally similar. Therefore, when the genes are embedded in the feature space, the genes of the same family should be closer and form gene clusters. However, a gene may belong to multiple gene families, and there may be overlap between gene families. In order to deal with such complex relationships, we use the gene2vec embedding method^23^ to list all genes with family co-belonging relationship as gene pairs when constructing training samples. Gene2vec can adaptively adjust the frequency of gene according to the size of the gene family and the number of families it belongs to, and fully consider the relationship between a single gene and other genes when it belongs to multiple families. In our study, 1645 gene families of humans and 1539 gene families of mice are used for embedding. The embedding of each gene is 768-dimensional.

#### Gene regulatory network (GRN) embeddings

GRN refers to the network formed by the interaction between genes in the cell or a specific genome. Among many interaction relationships, the regulatory relationship based on transcriptional expression between genes is particularly important. Generally, some genes can control the expression level of other genes through transcriptional expression. Usually, the GRN is a directed graph, each gene represents a node, and the control relationship between genes is described by weighted directed graphs in the GRN. Our goal is to refine the regulatory network as much as possible, based on the regulatory relationships that have been discovered and to learn more unknown interactions between genes. In different tissue cells, the regulatory relationship between genes is not static. Our goal is to make genes that have more regulatory relationships get closer in embedding space. Thus, we count the gene pairs with regulatory relations from GRN. These gene pairs are used for embedding through the gene2vec method^23^. The more the number of regulatory relations between gene pairs, the more they appear during training, and finally the closer they are after encoding. The embedding of each gene is 768-dimensional.

### Multiple downstream tasks

#### Single-species cell type annotation

The fine-tuning objective is identifying the type of each cell by utilizing its cell embedding generated by GeneCompass. Specifically, a fully connected layer is added to class token to predict the cell type from its cell embedding. The hyper-meters are as follows: epoch, 30; max learning rate, 5e–5; learning scheduler, linear with warmup; optimizer, Adamw with weight decay 0.001; warmup steps, 100; weight decay, 0.001; batch size, 10. The cross-entropy loss between the predicted cell type probabilities and ground truth is optimized as low as possible. For human-specific task, we benchmarked GeneCompass against the transformer-based method Geneformer on human multiple sclerosis (hMS), human lung (hLung), and human liver (hLiver) datasets. For mouse-specific task, the results GeneCompass with and without pre-training are compared on mouse brain (mBrain), mouse lung (mLung), and mouse pancreas (mPancreas) datasets. Meanwhile, we benchmarked GeneCompass against the method TOSICA on mouse pancreas (mPancreas) dataset. The details of datasets used for model training and validation can be found in Supplementary Material.

#### Cross-species cell type annotation

We aim to perform integration and cell-type assignment while preserving biological variability by utilizing the universal gene embeddings from our generative pre-trained model. We apply GeneCompass to CAME^28^ which is a heterogeneous graph neural network called GeneCompass-CAME, where cells and genes are modeled as heterogeneous nodes. Also, like CAME, we create the heterogeneous graph with six heterogeneous types: ‘cell to gene’, ‘gene to cell’, ‘cell to cell’, ‘gene to gene’, ‘cell self-loop’, ‘gene self-loop’, where we denote the corresponding weights (shared across species) as *W_cg_* , *W_gc_*, *W_cc_*, *W_gg_*, *W_c_* and *W_g_*, respectively. Unlike CAME, GeneCompass -CAME adopts gene embeddings from our generative pre-trained model as input.

Here, we denote a gene expression matrix with N cells and M genes as *X* ∈ *R^N^*^×*M*^, and the corresponding pre-trained cell embeddings as *X_e_* ∈ *R^N^*^×*P*^ . Taking the pre-trained cell embeddings as input, the initial embedding (the 0-th layer) for each cell *i* is calculated as:

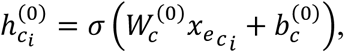

where σ is the leaky ReLU activation function with a negative slope of 0.05 and 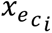 is the pre-trained embeddings for cell *i*, 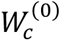 and 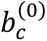 are learnable weight and bias vectors. Genes, unlike cells, lack of pre-trained embedding, so, we aggregate the pre-trained embeddings for corresponding neighbor cells as the initial embedding for each gene as follows:

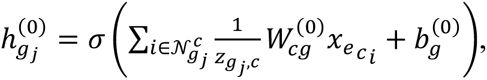

where 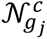 is the set of cells that have expressed the gene *j*, 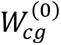 and 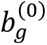 are learnable weight and bias vectors.

For the remaining layers, we use the same strategy as CAME, where we update the embeddings of each node under the guidance of the heterogeneous graph and use a graph attention layer with multi-head mechanism as the cell-type classifier. At last, we take cross-entropy loss with label smoothing mechanism^46^ as training loss and adopt Adam optimizer^47^ for training. Also, we use AMI to select the best checkpoint, which is also consistent with CAME. To evaluate the performance of cell-type assignment, we adopted accuracy as metrics which measures the proportion of correctly classified cells out of the total number of cells. The details of cross-species datasets used in our study can be found in Supplementary Material.

#### Gene regulatory network prediction

This task involves the prediction of interactions and relationships among genes to gain insights into how genes work together to control cellular processes. We employ the DeepSEM framework^29^ to evaluate the performance of inferring Gene Regulatory Networks (GRNs) with and without GeneCompass embedding. We use Immune Human dataset provided by scGPT^15^, and generate gene embedding from pre-trained GeneCompass model as input of a multivariate statistical model to analyze structural relationships among different random variables^48^. we calculate cosine similarity by following equation:

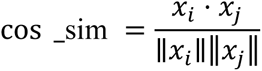

where x_i_ or x_j_ is a 2-dimensional vector belongs to gene embedding matrix generated from pre-trained GeneCompass with size ( n genes, embed size). ∥ x ∥ is the Euclidean norm of vector x.

For the implementation of GRN inference. Firstly, we feed the log-transformed scRNA-seq expression data after Z-normalizing into the neural network, the datasets come from BEELINE framework^49^, ground-truth GRNs are available from human embryonic stem cells (hESC)^50^, and the information source: non-specific ChIP-seq^51–53^, collected from the STRING database^54^. Secondly, we initialized MLPs by using the ‘kaiming_uniform’^4^ and initialized W by setting the matrix diagonal as zeros and the others following a Gaussian distribution N(1/(*m* − 1), ε^2^), in which *m* stands for number of genes and ε denotes a small value to avoid being trapped in the local optimal. The values on the diagonal are fixed as zeroes in the whole training process to guarantee that W can learn the regulatory network between genes. Finally, we add cosine similarity generated from GeneCompass as extra information to indicate the GRN learned by DeepSEM (Extended Fig. 2a).

#### Drug dose-response prediction

This task refers to the relationship between the drug dosage and the biological response in a cell. We employ the Compositional Perturbation Autoencoder (CPA)^30^ model to encode and learn transcriptional drug responses across different cell types, dose, and drug combinations, and evaluate the performance of predicting drug dose response with and without GeneCompass embedding. We use a dataset 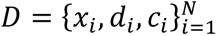 where each x_i_ ∈ *R*^G^ describes the gene expression of G genes from cell *i*. If d_id,i_ = 0, this means that perturbation j was not applied to cell *i*. Unless stated otherwise, the sequel assumes column vectors. Similarly, the vector of vectors c_i_ = (c_i.1_… ….c_i.K_) contains additional discrete covariates such as cell types or species, where each covariate is itself a vector. More specifically, c_i,j_ is a K_j_-dimensional one-hot vector. For the implementation of dose response, we train the CPA model using a dataset provided by Srivatsan et al.^55^, then we load the pre-trained CPA model, and implement CPA to reconstruct the expression response of compounds, with variable drug-dose combinations by following steps: (1) encoding the gene expression x_i_ into an estimated basal state 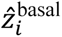 that does not contain any information about (d_i_, c_i_), (2) combining 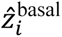 with learnable embeddings about (d_i_, c_i_), and we add embedding collected from model (Extended Fig. 2b).

#### Gene expression profile prediction

This task refers to estimate the levels of gene expression for a set of genes in a biological sample based on various biological conditions. We employ the DeepCE^31^ model designed for phenotype-based compound screening to evaluate the performance of GeneCompass embedding by adding it in DeepCE’s first component as extra information.

The DeepCE model consists of three components as follows: the first component includes the feature transformation component (GCN), pre-trained network (human protein–protein interaction network, extracted from STRING database^54^) and feed-forward neural network; the second component contains an interaction network (multi-head attention) and the last component has a prediction network (two-layer feed-forward neural network with a rectified linear unit activation function)^31^. DeepCE captures features from chemical compound, L1000 genes^56^, cell and dosage in the first component, and generates high-level feature associations (features from L1000 genes and chemical compound) in the second component. In the last component, all the features learned from the previous components will be concatenated as high-level features to predict gene expression profiles (Extended Fig. 2c).

For the implementation of predicting drug-induced gene expression perturbations, we add embedding collected from GeneCompass at the pre-trained network in DeepCE’s first component as extra information, then we run DeepCE model to test whether GeneCompass would benefit the performance of DeepCE or not.

#### Gene dosage sensitivity predictions

Distinguishing dosage-sensitive versus dosage-insensitive transcription factors is critical for explaining copy number variation (CNVs) in gene diagnosis. The traditional approach used conservation and allele frequency to predict dosage sensitivity. However, these characteristics do not vary with cell state and could not capture the specific tissues that would be influenced by dosage change of the gene. Following the protocol of Geneformer^16^, we use 10,000 random single-cell transcriptions to fine-tune GeneCompass to distinguish dosage-sensitive and dosage-insensitive transcription factors.

#### *In silico* perturbation

GEARS learned the gene embedding from the gene co-expression knowledge graph. This embedding is then combined with the perturbation embedding from the Gene Ontology (GO)-derived knowledge graph to predict the post-perturbation expression. In our study, we replaced GEARS’ gene embedding with GeneCompass’ gene embedding, utilizing the default parameter which set the length of gene embedding as 64. To handle cases where a gene, such as gene A, is not listed in the token list of a GeneCompass, we rank the genes and use the embedding of the gene that precedes gene A in the sorted list. We trained the GeneCompass model (epoch=10) using the Norman^57^ dataset provided by GEARS to predict gene expression after one-gene and two-gene perturbation. The loss function used for training is a mean squared error (MSE) for most differentially expressed genes, which was measured on the 20 genes with the largest difference between predicted expression and true post-perturbation expression.

We then evaluated GeneCompass’ performance on the perturbation of one or two genes whose data had been held out at the time of training, and thus those genes had not been seen experimentally perturbed during training. Analysis of experimental data showed that the expression of genes after perturbation is distributed around the mean of the dataset like a normal distribution. Therefore, we use the mean expression change of the truth experiment data as the standard for predicting accuracy. For the results of predicted expression, we calculate the absolute difference between the predicted values and the mean experiment expression values and select the top 20 genes with the most significant changes in the perturbed experiment to calculate the sum of the differences. This is defined as the top 20 DE deviation and serves as a criterion to evaluate the accuracy of predicted results for each perturbation, using the following formula.

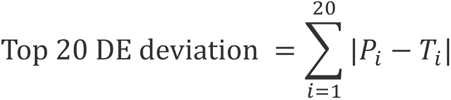

where *P*_i_ is the predicted postperturbation expression change by GeneCompass or GEARS for gene *i*, and *T*_i_ is the ground truth of mean expression change in the experiments.

#### *In silico* quantitative perturbation

The task is designed to simulate the cell reprogramming and differentiation inspired by Geneformer^16^. The perturbation state is characterized by the cell and gene embeddings. The *in silico* quantitative knock out is applied by decreasing the target gene’s expression value in the single-cell transcriptome input to GeneCompass. The *in silico* overexpression is realized by increasing the target gene’s expression value to a specific value. Note that our method could overexpress or decrease gene expressions to any value *in silico*. After gene expression manipulation, the single-cell transcriptome is reordered to simulate the cell state after perturbation. The post-perturbation embeddings are obtained by the forward propagation process of the encoder in GeneCompass. The effect of *in silico* quantitative perturbation is measured by calculating cosine similarity between the post-perturbation and ground-truth embeddings, which is defined as follows:

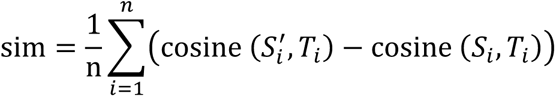

where *S_i_* denotes the source cell state and *T_i_* denotes the target cell states. The *in silico* perturbation state is denoted as 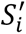

## Data and code availability

All primary data presented in this study will be deposited in a public database, and all codes will be uploaded to GitHub: https://github.com/xCompass-AI/GeneCompass.

## Acknowledgments

We would like to thank Dr. Dangsheng Li, Dr. Baoyang Hu and Dr. Wei Li for their invaluable comments and guidance. We thank Chuanyang Zhang, Jiajia Wang, Chengrui Wang, Yan Chen, Yang Wang, Qirui Gu, Shan Zong, Huimin He from the Computer Network Information Center, Chinese Academy of Sciences for their work in building website interface for model applications and model visualization. We thank Ran Zhang, Meng Xiao and the data process group from the X-Compass Consortium for their assistance in data collection and pre-processing. We are grateful to Dr. Weiwei Zhai, Dr. Liang Ma and Dr. Fei Gao from the Institute of Zoology for their comments in setting downstream tasks. We also appreciate Dr. Wei Yang, Xuewei Yuan and Jie Zhang for their outstanding management support. We thank Beijing Super Cloud Computing Center for their GPU resources for experiments. This work was also supported by CAS Project for Young Scientists in Basic Research, Grant No. YSBR-076.

## APPENDIX The X-Compass Consortium Members

### Institute of Zoology, Chinese Academy of Sciences

Xin Li, Hongmei Wang, Baoyang Hu, Wei Li, Fei Gao, Jingtao Guo, Leqian Yu, Qi Gu, Weiwei Zhai, Zhengting Zou, Guihai Feng, Wenhao Liu, Yao Tian, Chen Fang, Jingxi Dong, Yana Liu, Jingqi Yu, Wenhui Wu, Xinxin Lin, Cong Li, Yu Zou, Yongshun Ren, Fan Li, Yixiao Zhao, Yike Xin, Longfei Han, Shuyang Jiang, Kai Ma, Qicheng Chen, Haoyuan Wang, Huanhuan Wu, Chaofan He, Yilong Hu, Shuyu Guo, Yiyun Li

#### Computer Network Information Center, Chinese Academy of Sciences

Yuanchun Zhou, Yangang Wang, Xuezhi Wang, Pengfei Wang, Fei Li, Zhen Meng, Zheng Li, Zaitian Wang, Ping Xu, Wentao Cui, Zhilong Hu, Huimin He, Shan Zong, Jiajia Wang, Yan Chen, Chunyang Zhang, Chengrui Wang, Qingqing Long, Ran Zhang, Meng Xiao, Qinmeng Yang, Zijian Wang, Yining Wang

#### Institute of Computing Technology, Chinese Academy of Sciences

Yiqiang Chen, Yi Zhao, Xiaodong Yang, Dechao Bu, Xin Qin, Jiaxin Qin, Zhaohui Yang, Chenhao Li, Zhufeng Xu, Zeyuan Zhang, Xiaoning Qi, Shubai Chen, Wuliang Huang, Yaning Li

#### Institute of Automation, Chinese Academy of Sciences

Ge Yang, Jing Liu, Guole Liu, Jie Jiang, Xingjian He, Liqun Zhong, Yaoru Luo, Jiaheng Zhou, Zichen Wang, Qinxuan Luo, Ziwen Liu, Ao Li, Teng Wang, Yiming Huang, Handong Li

#### Academy of Mathematics and Systems Science, Chinese Academy of Sciences

Yong Wang, Shihua Zhang, Jiahao Zhang, Yiyang Zhang, Shirui Li, Zhongming Liang, Zhenpeng Man, Kangning Dong, Qunlun Shen

---

**Extended Fig.1.**
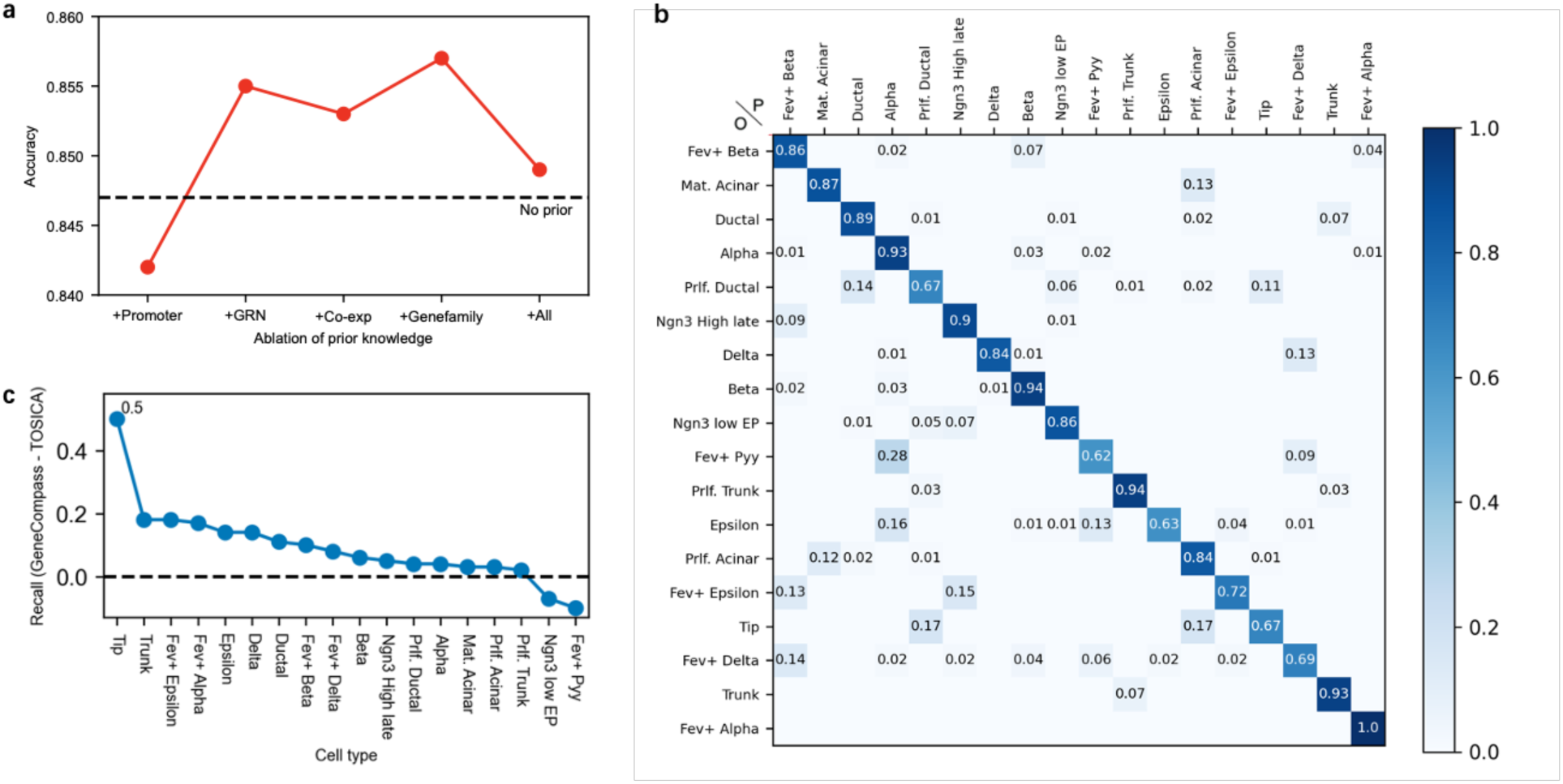
Prior knowledge ablation and results of cell type annotation. **a,** The ablation study of four kinds of prior knowledge of GeneCompass on hMS dataset, where ‘+All’ means all prior knowledges are concatenated. **b**, The confusion matrix of GeneCompass on mPancreas dataset. **c**, The difference of recall between GeneCompass and TOSICA for different cell types on mPancreas dataset.

**Extended Fig.2.**
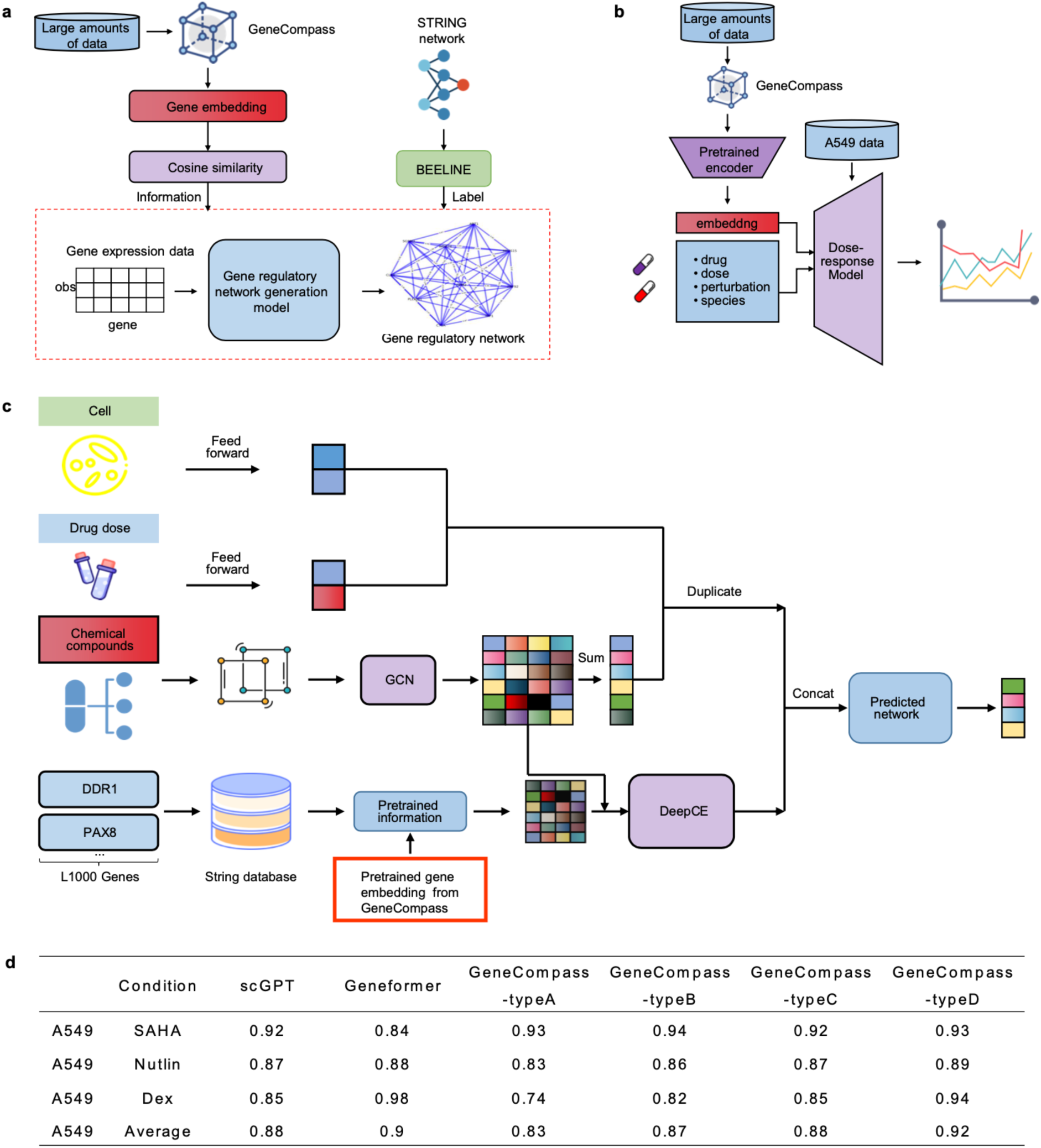
Workflows of three downstream tasks, i.e., GRN inference (a), drug response (b), and gene expression profile (c) prediction. **d,** The results of dose response prediction task, Type A, B, C and D denote results of GeneCompass pre-trained by 0.05, 0.5, 5 and 55 million human datasets, respectively.

