## Supplementary figures and images for "GeneCompass: Deciphering Universal Gene Regulatory Mechanisms with Knowledge-Informed Cross-Species Foundation Model"

### Fig S1

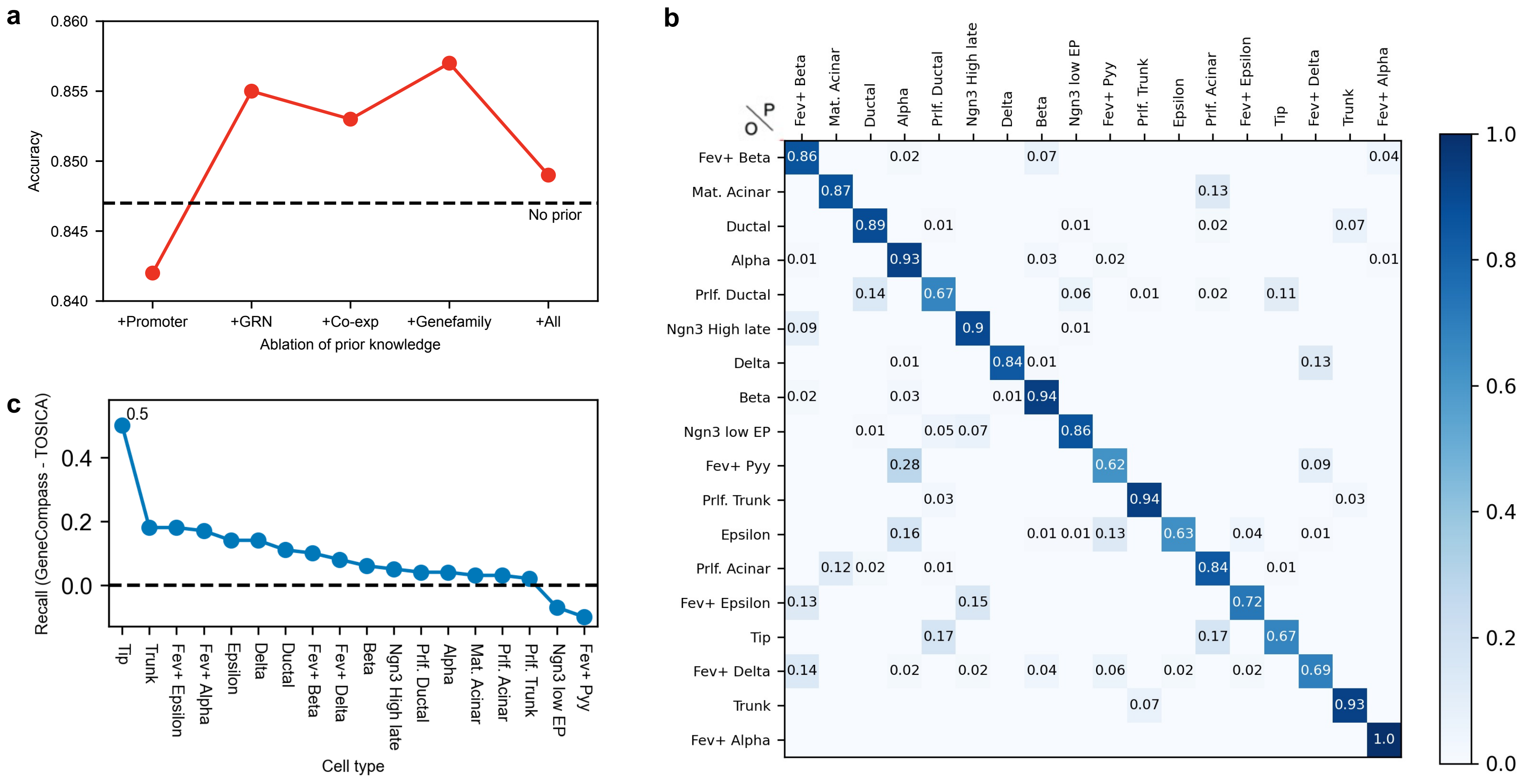
