## Supplementary material for "GeneCompass: Deciphering Universal Gene Regulatory Mechanisms with Knowledge-Informed Cross-Species Foundation Model": Fig S2

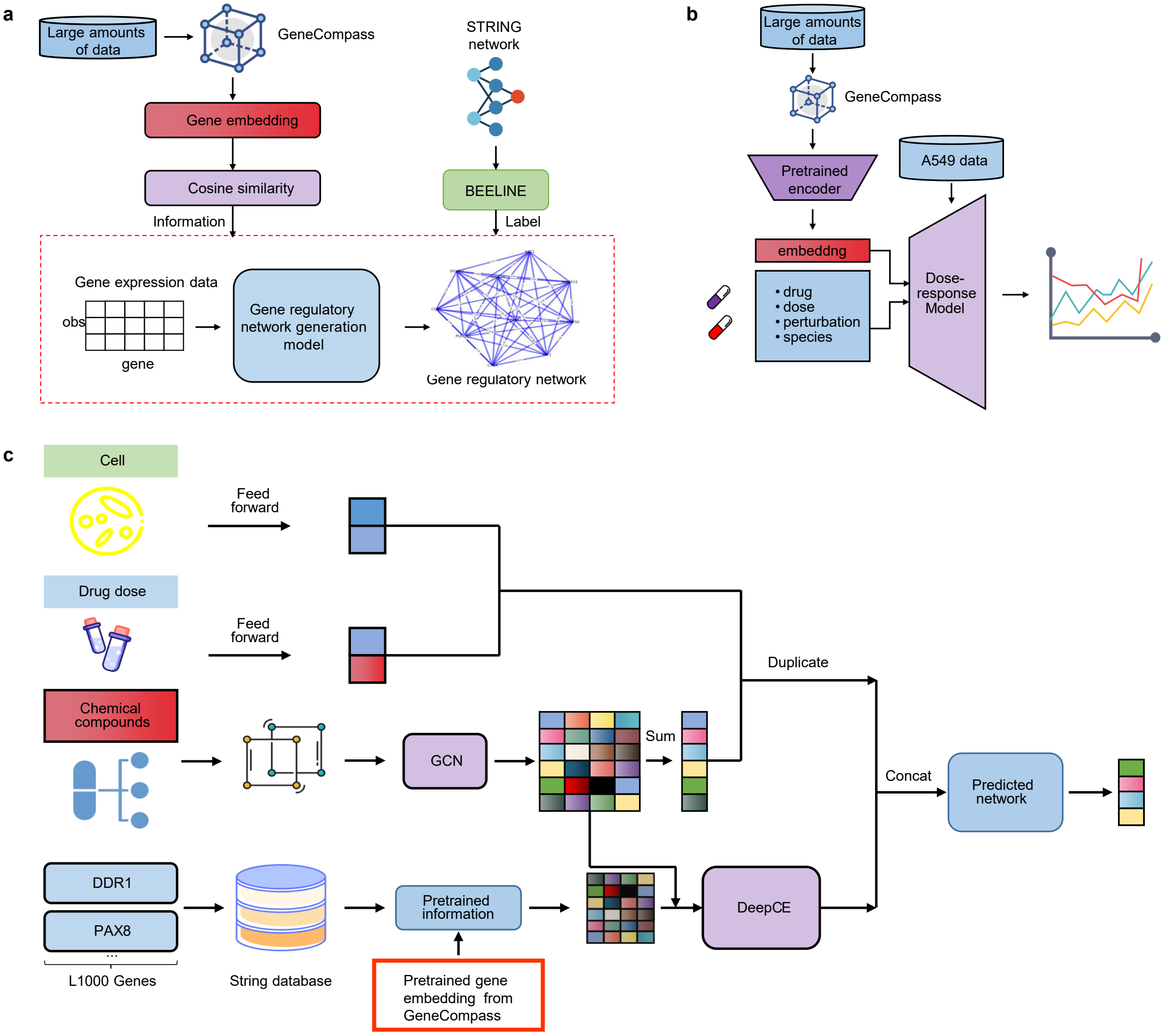


**d**

| | Condition | scGPT | Geneformer | GeneCompass -typeA | GeneCompass -typeB | GeneCompass -typeC | GeneCompass -typeD |
|---|---|---|---|---|---|---|---|
| A549 | SAHA | 0.92 | 0.84 | 0.93 | 0.94 | 0.92 | 0.93 |
| A549 | Nutlin | 0.87 | 0.88 | 0.83 | 0.86 | 0.87 | 0.89 |
| A549 | Dex | 0.85 | 0.98 | 0.74 | 0.82 | 0.85 | 0.94 |
| A549 | Average | 0.88 | 0.9 | 0.83 | 0.87 | 0.88 | 0.92 |
